# Neuronal miR-17-5p contributes to interhemispheric cortical connectivity defects induced by prenatal alcohol exposure

**DOI:** 10.1101/2022.12.30.522325

**Authors:** Mike Altounian, Anaïs Bellon, Fanny Mann

**Affiliations:** Aix Marseille Univ, CNRS, IBDM, Marseille, France; Aix Marseille Univ, INSERM, INMED, Marseille, France

**Keywords:** Prenatal alcohol exposure, Establishment of brain connectivity, Axon guidance, miRNA, Contralateral targeting, Somatosensory and motor cortex, In utero electroporation, 3D imaging

## Abstract

Prenatal alcohol exposure (PAE) is the leading cause of non-genetic intellectual disabilities in the Western world and is responsible of a wide spectrum of neurodevelopmental disorders referred to as Fetal Alcohol Spectrum Disorders (FASD). Structural and functional deficits in brain connectivity have been reported in FASD patients; still, whether and how PAE affects the axonal development of neurons and disrupts the wiring between brain regions is not known. Here, we developed a mouse model of moderate alcohol exposure during prenatal brain wiring to study the impact of PAE on corpus callosum (CC) development, a major white matter tract reported to be affected in FASD patients. Our results show that PAE induces aberrant navigation of interhemispheric CC axons that persist even after the end of the exposure, causing their ectopic termination in the contralateral cortex. Furthermore, these defects in interhemispheric connectivity persist into adulthood and are associated with defective bilateral sensorimotor coordination in behavioral tasks requiring cortical control and interhemispheric communication. Finally, we identified neuronal miR-17-5p and its target Ephrin type A receptor 4 (EphA4) as mediators of the effect of alcohol on the contralateral targeting of CC axons. Taken together, our results suggest that alteration of miRNA-mediated regulation of axon guidance signaling by prenatal alcohol exposure affects interhemispheric cortical connectivity and associated behavior in FASD.

## Introduction

The effects of the environment and lifestyle on human health have become a major concern in our societies. The developing fetal brain is particularly vulnerable to external insults that can lead to long-term neurological disorders. Among these external factors, maternal consumption of alcohol during pregnancy causes a wide spectrum of neurodevelopmental disorders referred to as Fetal Alcohol Spectrum Disorders (FASD) and is the leading cause of non-genetic intellectual disabilities in the Western world, with a prevalence of approximately 9 on 1000 births (1). Indeed, prenatal alcohol exposure (PAE) disrupts brain development at any point in gestation, causing lifelong cognitive, behavioral, emotional, and adaptive functioning deficits. Deficits vary from mild to severe depending of the time of exposure, the amount of alcohol consumed and the drinking patterns (2). Research based on animal models of FASD have shown that multiple steps in neuronal development can be affected by alcohol: alterations in the proliferation and survival of neural progenitors, differentiation, migration and survival of postmitotic neurons, synapse genesis and function have been reported (3). Despite these advances, it is still not known whether and how PAE affects the axonal development of neurons and disrupts the wiring between brain regions that underlies our behavior.

Structural imaging studies have reported reduced volume and disorganization of white matter tracts in FASD children and adults (4,5). Functional connectivity studies also revealed that FASD patients present reduced inter-network functional connectivity (circuit between remote region) while their intra-network functional connectivity (circuit inside a defined region) is unchanged or even increased compared to controls, suggesting underlying structural deficits of long range brain connectivity (6,7). One particular target of alcohol neuroteratogenicity is the corpus callosum (CC), which is the largest fiber tract connecting the left and right cortical hemispheres. Structural Magnetic Resonance Imaging (MRI) studies of children and adults with prenatal exposure to alcohol have revealed a continuum of CC abnormalities ranging from complete absence (agenesis), underdevelopment (hypoplasia) and spatial displacement (8,9). These variations in the CC may underlie the impaired interhemispheric communication that has been reported in FASD, particularly in visual-motor integration and bimanual coordination (10).

Abnormalities in interhemispheric brain connectivity in individuals prenatally exposed to alcohol may be the repercussions of the pleiotropic effects of alcohol on earlier stages of embryonic brain development (e.g. neuronal production or migration), or may result from a direct impact of alcohol on the navigation of neuronal axons to reach their final destination. *In vitro* evidence suggests that developing axons could be a direct target for alcohol. Alcohol treatment in primary mouse neuron cultures delays axonogenesis (11), disrupts the structure and motility of axonal growth cones (12,13)and affects the way axons respond to molecular cues, such as the cell adhesion molecule L1CAM, the neurotrophic factor BDNF, and the axon guidance molecules Semaphorin 3A and Netrin-1 (14,15). How these in vitro data relate to the in vivo effects of alcohol on neural circuit formation is not known.

Although the devastating effects of PAE have long been known, our knowledge on the underlying causative and protective mechanisms has just started to emerge. Accumulating evidence suggests a predominant contribution of epigenetic mechanisms such as DNA methylation and histones modifications(16). Recently, studies have also reported that developmental exposure to alcohol alters the expression of another type of epigenetic regulators: microRNAs (miRNAs) (17), a class of endogenously expressed small noncoding RNAs that regulate the expression of target genes at the post-transcriptional level. While miRNAs constitute biomarkers for exposure to environmental factors and are involved in diverse pathophysiological processes (18), the role of alcohol-sensitive miRNAs in the pathogenesis of FASD remains largely unknown. Interestingly, miRNA have emerged as key regulators of axonal development and brain connectivity (19). Aberrant development of several axon tract, including brain commissures, has been reported after the knockdown of canonical miRNAs in Dicer mutant mice (20). In addition, several studies have shown that miRNAs can act in different cellular compartments to modulate axon navigation (21). MiRNAs can act in the neuronal cell body to regulate the level and timing of expression of genes required for proper axon growth and navigation (22). In addition, selected populations of miRNAs can be transported and stored in developing axons where they directly modulate the local translation of genes involved in axon metabolism, growth, branching and guidance in response to external chemical cues (23–26). Thus, miRNA are potential candidates to mediate the effects of alcohol on the development of brain circuits.

Here, we developed a mouse model of alcohol exposure during prenatal brain wiring to study the effect of alcohol on CC development and long-term functional consequences. Our results showed that alcohol induces aberrant axonal navigation that persists even after cessation of exposure, causing ectopic termination of CC axons in the contralateral cortex. We also identified miR-17-5p and its target EphA4 as mediators of the effect of alcohol on the contralateral targeting of CC axons. Taken together, our results provide insight into the mechanisms by which PAE through miRNA regulation affects interhemispheric cortical connectivity in FASD.

## Methods

### Animals

Animal procedures were conducted in accordance with the guidelines of the French Ministry of Agriculture (approval number F1305521) and approved by the local ethics committee (CE14 approval number APAFIS# 20223-2021011909315100 v1). All experiments were carried out with Swiss mice (RjOrl:SWISS) purchased from Janvier Labs.

### Alcohol administration and dosage

Alcohol was administered to pregnant mice by daily intraperitoneal injection of 25% EtOH (in saline (0.9% NaCl), final 2.0 g/kg body weight) or saline solution (control) from E15.5 to E18.5. Blood, amniotic fluid and fetal brain alcohol concentrations were mesurted using the Ethanol Assay Kit (Cat. #ab272531; Abcam, Cambridge, UK).

### In Utero Electroporation

In utero electroporation was performed at E15.5 as described in (27). Briefly, pregnant mice were anesthetized. A laparotomy was performed to expose the uterine horns. 0.5-1µl of plasmid DNA solution was injected in a lateral ventricle of E15.5 mouse brain, and electroporated into the developing sensorimotor cortex. Uterine horns were then placed back in the abdominal cavity and the abdominal muscles and skin were sutured. Electroporation of 1µg/µl of pCAGGS-IRES-tdTomato reporter plasmid was performed to trace electroporated axons, together with 2 µg/µl of miR-17-5 sponge plasmid or control plasmid when required.

### Retrograde tracing of callosal projections

Adult (8–9 weeks) control or PAE mice were anaesthetized, and placed in a stereotaxic apparatus. 200uL of Cholera toxin subunit B (CTB) conjugated to Alexa Fluor 555 (Life technologies, Cat# C34776) or Alexa Fluor 647 (Life technologies, Cat# C34778) (2 μg/μL in PBS) were injected in S1 and M1 respectively, with a nanolitre injector (Nanoject III, Drummond, Cat# 3-000-207) according to coordinates described in table 2. After surgery, mice were injected with 5 mg/kg carprofen (Rimadyl, Pfizer).

### Axonal length measure

The 10 longest axons in the contralateral hemisphere were traced from the midline using FIJI NeuronJ plugin, and the length mesured (28).

### Contralateral innervation

Quantification of contralateral innervation ratio was performed by measuring the size of the contralateral projection area (from the position of the first axon observed from the midline in the contralateral cortex, to the farthest). Values were normalized to the corresponding ipsilateral electroporated area (from the most lateral to the most medial neuron electroporated). To analyze the innervation in each cortical area, a threshold was applied to remove the background and ROI corresponding to the different cortical area were traced in Fiji-ImageJ (29). The density of tdtomato signal was measured for each ROI. Data were normalized to the density of ipsilateral signal to avoid any differences due to electroporation efficiency.

### MicroRNA Sponge

miR-17-5p sponge sequence is composed of six copies of miR-17-5p binding sites (MBS) ctacctgcactCCCgcactttg (bulged sequence in capitals). MBS are separated by diverse spacer sequences (GACG, GTTAT or GGAA). To assess the in vivo function of miR-17-5p in CPNs, the sequence containing 6MBS for miR-17-5p was inserted into pNeuroD-IRES-GFP vector (30) (gift from L. Nguyen, C.H.U, Liège, Belgium) using XhoI and SalI.

### Cell dissociation and FACS

Electroporated cortical regions of P0 mice brains were dissected out based on the observation of tdtomato fluorescence under a binocular microscope (Leica MZFLIII). Dissected cortices were subjected to cell dissociation using a papain solution and mechanical dissociation as described previously (31). Single cell suspension was loaded in FACSAriaII (BD Biosciences, San Jose, USA) and appropriate gates (Supplemental figure 8), were set based on relative levels of tdTomato to isolate electroporated CPN using FACSDIVA version 8.0.1 flow cytometry gating software. FACS-sorted cells (≈ 98% tdTomato+ cells) were harvested in RLT and RNA extracted with RNeasy Micro kit (Cat. # Qiagen). Around 100 000 tdTomato+ cells were sorted from 10 electroporated-brains for each sample to sequence. Around 50ng of RNA were extract for each sample (mean RNA integrity number of 9,32 for control condition and 9,12 for sponge-17 condition).

#### RT-qPCR of miRNAs were performed using

miRCURY LNA RT kit (Cat. #339340) and miRCURY LNA SYBR® Green PCR Kit. RT-qPCR of mRNAs were performed using SuperScript III Reverse Transcriptase (Cat. #18080-044, Invitrogen) and PowerUp SYBR® Green master mix (Cat. # A25742, Thermofisher scientific). Please refer to supplementary methods for details and probe sequences.

### Quantification of the distribution of CTB retrolabeled neurons

Seven days after CTB injection, mice were anaesthetized and perfused intracardially with 4% PFA. Brains were dissected, post-fixed in 4% PFA overnight at 4 °C and 100um thick coronal vibratome sections were made (Leica VT1000S). Images of both ipsilateral (injected) cortex and contralateral (retrolabeled) cortex were acquired with an Apotome imager M2 (ZEISS). A bin system was established along the mediolateral axis of the contralateral cortex, with each bin corresponding to a 250um wide column. Bin 4 was centered on the homotopic region to M1 injection site. The number of Alexa Fluor 555+ and Alexa Fluor 647+ neurons was counted in each bin.

### Rotarod assay

Control or PAE P60 Mice were placed on a rod that slowly accelerated from 4 rpm to 44 rpm over 5 min (rotarod apparatus, LSI Letica Scientific Instruments) and the latency to fall off during this period was recorded. The test was done on 4 consecutive days. Each day, the animals were tested three times separated by at least 5 min resting period. Data presented were acquired on the last day.

### Complex wheel-running assay and tamapin injection

P60 Mice were housed in individual cages equipped with activity wheel and optical sensor to detect the number of wheel revolutions per time interval (Cat. # 80821S; Lafayette Instruments, Lafayette, USA). Animals were first allowed voluntary running on a training wheel with all 38 rungs intact to get accustomed to the task. After 10 days, wheels were replaced by complex wheels with 16 rungs missing according to the pattern shown in Fig. 3B and wheel-running activity was recorded for one week. Mice were then treated with tamapin (Cat. # 10TAM001-00100; Smartox Biotechnology, Saint-Égrève, France) administered at 15 µg/kg body weight via intraperitoneal injection every other day, or with control (saline) solution, and wheel-running activity was recorded for a week. Wheel revolutions were monitored with 1min intervals using Scurry Activity Monitoring Software (Lafayette Instruments, Cat. # 80821S). Data obtained during the dark phase were analyzed. Total distance run, maximum velocity among daily runs, number of runs and duration of run were calculated. 15 PAE mice and 15 sex-matched control mice were tested. Among them 10 mice per group were treated with tamapin. All mice except one control, excluded from the analysis, showed spontaneous running activity.

## Results

### Establishment of a murine model to study the effect of PAE on CC development

To assess the impact of PAE on the development of the CC, we established a mouse model in which embryos were exposed to alcohol during the initial phase of callosal axon growth, from embryonic day (E) 15.5 to E18.5 (Fig. 1A and 1B). During this period, the previously separate medial walls of the cerebral hemispheres begin to fuse, and a transient bridge-like or “sling” structure composed of glial cells and neurons forms at the midline. The first developing callosal axons from the cingulate cortex use this sling to cross the midline at E16.5, pioneering the path for later-developing neocortical callosal projection neurons (CPNs), which begin to cross a day later (32). PAE can have a profound impact on neural progenitor cell proliferation, which can result in the creation of fewer neurons and glial cells, reduced brain volumes (microcephaly), and gross morphological abnormalities of brain midline structures (33,34). Here, to minimize these effects, pregnant Swiss mice were treated intraperitoneally with subthreshold levels of EtOH (2 g/kg body weight) that caused no significant cortical abnormalities in other studies (35). Dosage of maternal blood alcohol concentration (BAC) showed that alcohol level raised rapidly, reaching a peak around 160 mg/dL 30 minutes after the injection and then gradually returning to normal 4 hours later (Fig. 1C). Such BACs are considered low to moderate in mice (34). To further assess the exposure of embryos to alcohol, we also measured EtOH concentration in the amniotic fluid and in the brain of the embryos. While the alcohol concentration in the amniotic fluid was comparable to maternal BAC, the concentration in the embryonic brain was lower with a maximum peak at 100 mg/dL. Thus, the PAE paradigm used here corresponds to a 4-hour daily challenge of the developing brain with alcohol concentrations < 100 mg/dL over a 4-day period before birth. Finally, we confirmed that this pattern of exposition did not induce major structural anomalies, since brain size, general morphology and cortical thickness were not affected at birth compared to vehicle-treated controls (Fig. 1D-1G).

**Figure 1:**
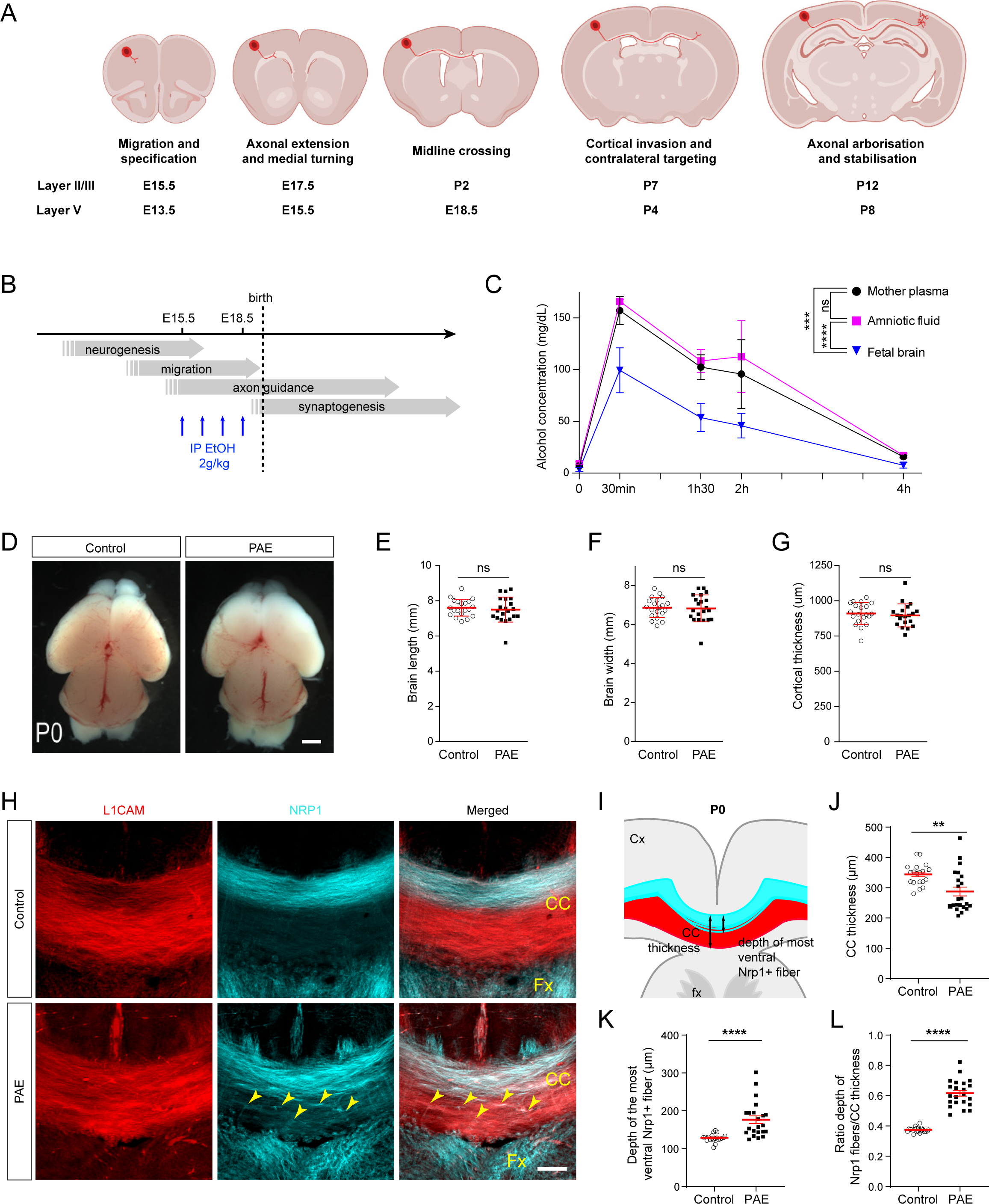
Moderate prenatal alcohol exposure affects the size and topographic organization of the callosal tract at birth. **A**, Developmental timeline of callosal projection development in the mouse. **B**, Schematic of EtOH injections. **C**, Evolution of alcohol concentration in maternal plasma, amniotic fluid and fetal brain after EtOH injection (time 0). Data are mean ± SEM, maternal blood: n= 3 pregnant control-injected and n=3 pregnant EtOH-injected mice ; amniotic fluid and fetal brain: n= 6 control and 6 PAE embryos from 3 litters. Maternal plasma vs amniotic fluid p= 0.3449 (Two-way ANOVA), maternal plasma vs Fetal brain p= 0.0006 (Two-way ANOVA), Amniotic fluid vs Fetal brain p= <0.0001 (Two-way ANOVA). **D**, Representative images of P0 mouse brains prenatally exposed to alcohol (PAE) or saline solution (control). **E-G**, Quantification of brain length (**E**) and width (**F**), and cortical thickness (**G**) in PAE and control P0 brains. Data are mean ± SEM, **E-F**, n=20 control and 21 PAE mice from 3 litters, **G**, n=10 control mice from 3 litters, 20 sections, and n=11 PAE mice from 3 litters, 20 sections. **E,** p= 0.5396 (Mann-Whitney test); **F**, p= 0.9621 (Mann-Whitney test); **G**, p=0.3802 (Mann-Whitney test). **H**, Immunostaining of L1CAM and Neuropilin 1 (NRP1) on tissue sections through the CC of PEA and control mice at P0. Arrowheads indicate ectopic Nrp1+ axons in the ventral part of the CC. **I**, Schematic illustrating measurement methods for analysis of CC thickness and position (depth) of the most ventral Nrp1^+^ fiber. **J-L**, Quantification of CC thickness (**J**), depth of the most dorsal Nrp1+ fiber (**K**) and the ratio depth of the most ventral Nrp1^+^ fiber /CC thickness (**L**) in PEA and control brains at P0. Data are mean ± SEM, n=15 control mice from 4 litters, 20 sections, and n=15 PAE mice from 4 litters, 22 sections. **J,** p= 0.0020 (unpaired t-test); **K**, p<0.0001 (Man –Whitney test); **L**, p<0.0001 (unpaired t-test). CC, corpus callosum; Fx, fornix. Scale bars: 500µm (**D**), 100µm (**H**)

### Prenatal exposure to EtOH induces CC midline defects

To determine whether early CC development was affected by the PAE paradigm described above, we first examined the midline of the CC at postnatal day (P) 0, approximately 24 h after the last EtOH injection. L1CAM immunostaining of coronal brain sections revealed a small but significant reduction in the thickness of the CC (Figure 1H-1J) in brains prenatally exposed to EtOH as compared to controls. This hypoplasia could indicate a defect in the production or specification of CPNs. However, immunostaining for key transcription factors expressed in early cortical projection neurons indicated no differences in the number and laminar distribution of the major neuronal subtypes, including SATB Homeobox 2 (Satb2)-expressing CPNs (36–38), between exposed and control cortices (Supplementary Fig. 1). CC defects may also be associated with reduced or disorganized guidepost cells. However, at P0, midline glial populations and subcallosal sling neurons were present and localized normally in brains exposed to alcohol (Supplementary Fig. 2A and 2B). Thus, it is likely that the observed reduction in CC may result from a delay in the extension of callosal axons across the midline, a process that is not yet complete at birth and continues into the first postnatal week (39).

In the developing mouse, callosal axons from the medial and lateral regions of the cerebral cortex are topographically segregated and project through the dorsal and ventral parts of the CC, respectively (40–42). This dorso-ventral arrangement is clearly visible in control mice where immunostaining with the surface receptor Neuropilin-1 (Nrp1) labels axons from the medial cortex in the dorsal half of the CC (Fig. 1H). However, this segregation appeared disrupted in P0 brains after subthreshold EtOH challenge, with some Nrp1^+^ axons defasciculating from the dorsal tract and invading the ventral portion of the CC from which they were normally excluded (Fig. 1H, 1K and 1L). A previous study reported abundant microglia in the CC and showed that activation of microglia caused by maternal inflammation resulted in a similar phenotype in the dorsoventral organization of the callosal tract (43). We thus evaluated the effects of subthreshold EtOH challenge on microglia *in vivo* by staining P0 brain sections with ionized calcium-binding adaptor protein-1 (Iba-1). We did not observe any difference in the localisation and number of microglial cells in the CC of brains exposed to EtOH (Supplementary Fig. 2C and 2D). Furthermore, co-staining with the reactive microglia markers CD68 and cyclooxygenase-2 (Cox2) showed no differences from control neonates, indicating that our PAE paradigm did not trigger classical activation of microglia (Supplementary Fig. 2C, 2E and 2F). Taken together, our data suggest that PAE affects early stages of CC axon growth across the midline, independent of major effects on midline guidepost cell populations.

### Prenatal exposure to EtOH alters postnatal CC development

The alterations in CC described in newborn mice could have an impact on later stages of CC development, despite the cessation of EtOH treatment after birth. Indeed, it has been proposed that the dorso-ventral position of axons in the CC, rather than their cortical origin, determines their contralateral projection after midline crossing (42). Given the altered dorso-ventral organization of the CC observed at birth, we analysed whether PAE alters the final targeting of callosal axons. To trace callosal projections, we labelled upper layer CPNs of the somatosensory cortex by *in utero* electroporation with a dtTomato reporter plasmid at E15.5, challenged the embryos with EtOH or vehicle and analysed final targeting of labelled axons at P8, the stage at which callosal projections have invaded and begun to arborize in the contralateral cortex. We first verified the dorso-ventral position of labelled axons at the midline. Labelled somatosensory axons travelled through the ventral CC as expected in controls, but were found both in the ventral and the dorsal CC in PAE individuals (Supplementary Figure 4A). We then measured the area targeted by electroporated axons in the contralateral cortex (Fig. 2A). To exclude effects caused by embryo-to-embryo variation in the electroporation sites, the size of the contralateral target area was normalized to the size of the ipsilateral electroporated area. Results showed an increase in the contralateral projection area in the EtOH exposed group compared to controls (Figure 2B-2D). Specifically, in control animals, labelled CPNs established projections to the contralateral somatosensory cortex that distributed with moderate density in the primary (S1) cortex and higher density at the primary/secondary (S1/S2) border region, consistent with previous findings (44) (Fig. 2C, 2E). In PAE brains, this innervation pattern was still visible but some electroporated axons ectopically invaded the adjacent cingulate and motor cortices (Fig. 2D and 2F-2L). At the same stage, analysis of early cortical areal patterning in PAE brains showed normal positioning of the boundary between somatosensory and motor cortex (Fig. 20-2Q). Thus, PAE disrupts not only the early phase of callosal axon growth at the midline, but also the establishment of contralateral callosal projections that occurs during the first weeks of postnatal development, when the brain was no longer exposed to alcohol.

**Figure 2:**
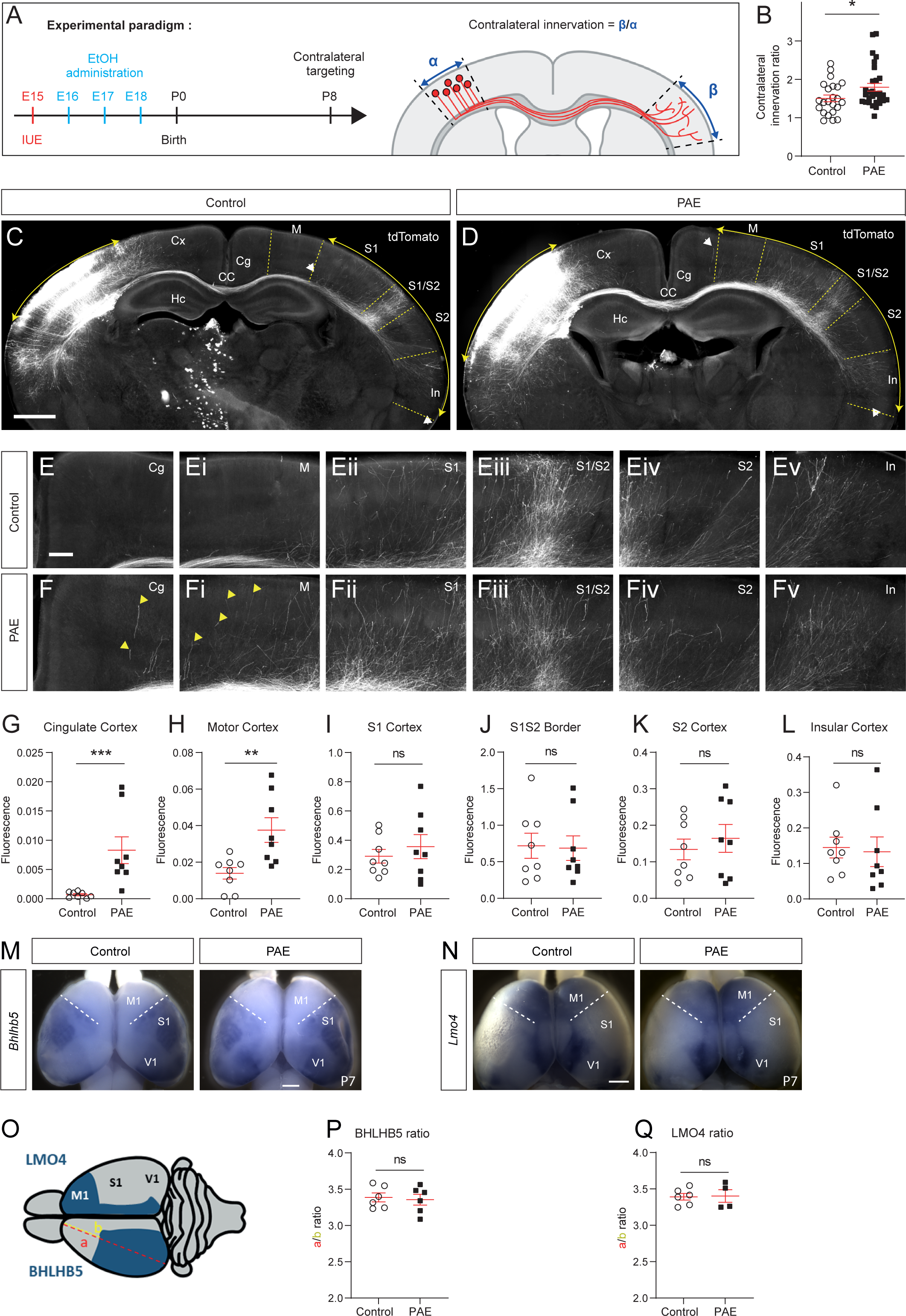
Abnormal heterotopic contralateral targeting of callosal projections in brains prenatally exposed to alcohol. **A,** Schematic of the experimental paradigm and quantification method of the contralateral innervation region. IUE: *in utero* electroporation. **B**, Quantification of the contralateral innervation in PEA and control brains at P8. Data are mean ± SEM, n=5 control mice from 2 litters, 23 sections, and n=6 PAE mice from 2 litters, 29 sections. p= 0.0485 (Mann Whitney test). **C** and **D**, Sections of P8 brains from control and PEA mice showing layer 2/3 callosal neurons of the somatosensory cortex labeled by in utero electroporation at E15.5 with a tdTomato expressing plasmid, and their projections into the contralateral cortical hemisphere. **E** and **F**, Higher resolution images of the different cortical regions shown in **C** and **D**, revealing ectopic callosal projections in the cingulate and motor cortex of PEA brains (yellow arrowheads). **G-L**, Quantification of tdTomato fluorescence in the different cortical areas of PEA and control brains. Data are mean ± SEM, n=8 control mice from 3 litters, 8 sections, and n=8 PAE mice from 3 litters, 8 sections. **G**, p= 0.0003 (Mann-Whitney test); **H**, p= 0.0047 (Mann-Whitney test); **I**, p= 0.7984 (Mann-Whitney test); **J**, p= 0.8785 (Mann-Whitney test); **K**, p= 0.6454 (Mann Whitney test); **L**, p= 0.5737 (Mann Whitney test). **M** and **N,** In situ hybridization for the regional markers *Bhlhb5* and *Lmo4* on whole brains of PAE and control mice at P7. **O,** Schematic representation of the expression territories of Bhlb5 and Lmo4 in mice brains at P7 and the analysis method. **P** and **Q**, Quantification of the rostro-caudal boundary defined by Bhlb5 and Lmo4 expression patterns in PEA and control brains at P7. Data are mean ± SEM ; **P,** n=6 control and 6 PAE mice from 2 litters, p= 0,8182 (Mann-Whitney test), **Q,** n=6 control and 4 PAE mice from 2 litters, p= 0,9143 (Mann-Whitney test). CC, corpus callosum; Cg, cingulate cortex; Cx, cortex; Hc, hippocampus, In, insular cortex; M, motor cortex; S1, primary somatosensory cortex; S2, secondary somatosensory cortex ; V1, primary visual cortex. Scale bars: 1200 µm (C, D), 1200 µm (Ci, Di), 300 µm (E, F), 150 µm (M, N).

### Prenatal exposure to EtOH impairs adult callosal interhemispheric connectivity

To determine whether the callosal axon targeting defect observed at P8 persists into adulthood, we analyzed the brain of adult (P60) mice prenatally exposed to EtOH. Adult PAE brains showed normal size and cortical thickness. The CC, which appeared hypotrophic at birth, had recovered a normal thickness and its myelination was similar to that of control brains (Supplementary Fig. 5). The pattern of interhemispheric connections was then analysed using retrograde fluorescent labelling with Alexa Fluor (AF) conjugates of CTB (cholera toxin B subunit). CTB-AF647 was injected into the primary motor cortex (M1) and CTB-AF555 into S1 and the mediolateral distribution of retrolabelled neuronal cell bodies in the contralateral cortex were analyzed (Fig. 3A and Supplementary Fig. 6). In control animals, 98.5% of CPNs retrolabelled from M1 were localized in the contralateral M1 (bins 1-7), whereas 98.8% of CPNs retrolabelled from S1 were distributed in the contralateral S1 cortex (bins 8-16), reflecting the homotopic nature of CC connectivity (Fig. 3B and 3C). However, in adult PEA mice, the distribution curves of CTB-AF647 and CTB-AF555 labelled neurons were flattened, with a significant decrease in CTB-AF647^+^ cells in bin 4 and 5 and CTB-AF555^+^ cells in bin 11, where the maximal peaks of cells were detected in control cortices. In addition, in contrast with controls, the distribution of CTB-AF647 and CTB-AF555 retrolabelled neurons overlapped, with the notable presence in the M1 cortex of CPNs retrolabelled from the contralateral S1 area (Fig. 3B and 3C). These results reveal that the topographic map of cortical interhemispheric connectivity is durably affected in adult mice that were exposed to EtOH *in utero*, with a decrease in homotypic projections and abnormal contralateral heterotopic projections between S1 and M1 areas.

**Figure 3:**
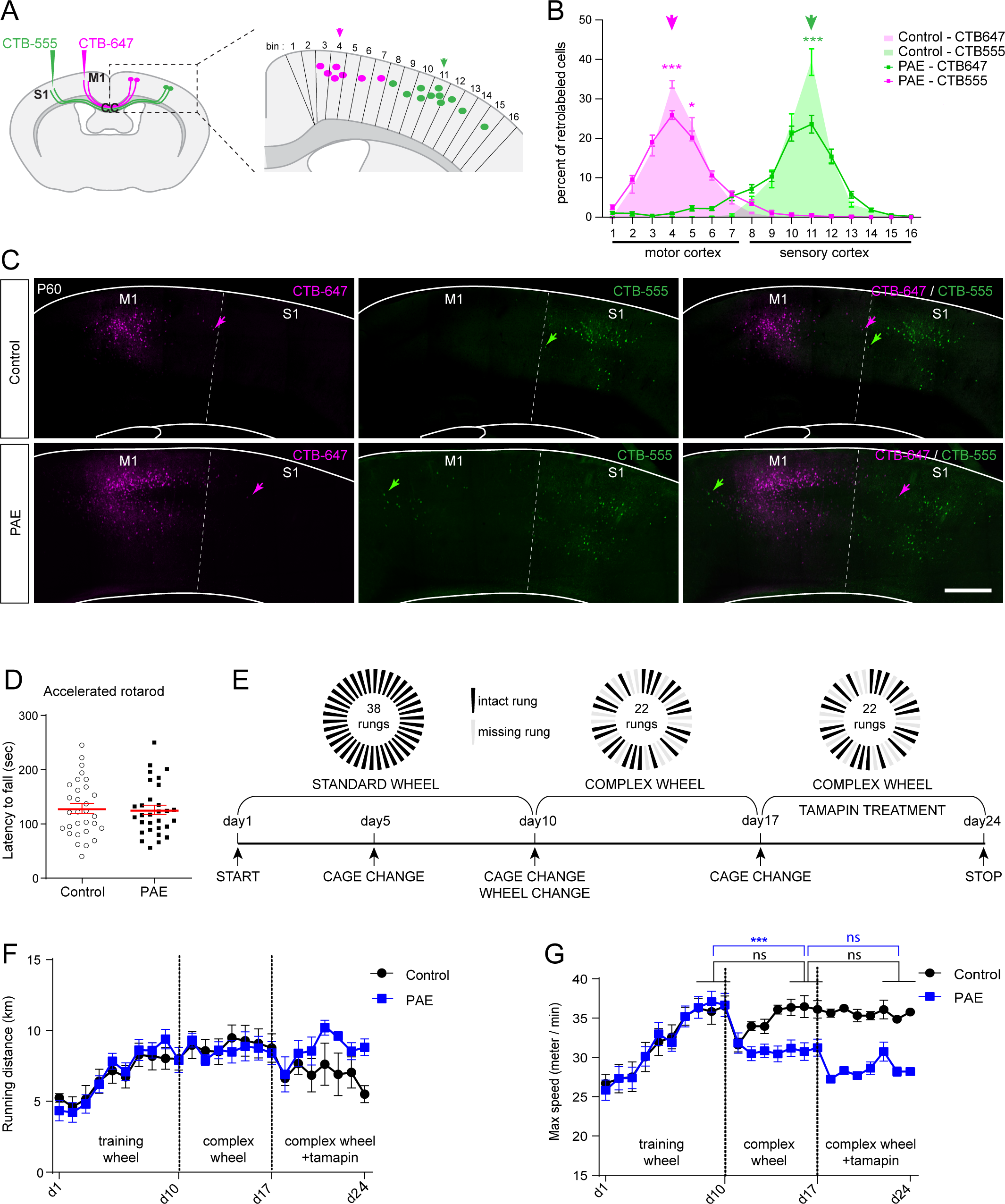
Contralateral CC connectivity defects are maintained in adult mice and correlate with long-term effect of PAE on bilateral sensorimotor coordination. **A,** Schematic of the experimental paradigm and analysis. Motor and somatosensory cortices of control and PEA brains were injected at P60 with CTB-AF647 (magenta) in M1 and CTB-AF555 (green) in S1. The distribution of CTB-AF647 ^+^ neurons and CTB-AF555 ^+^ neurons was quantified in the contralateral cortex segmented into 16 equidistant bins. Bins 1 to 7 correspond to the motor cortex and bins 1-16 to the somatosensory cortex. Magenta and green arrows point to the cortical regions homotopic to the injection sites of CTB-AF647 and CTB-AF555 respectively. **B**, Quantification of the distribution of CTB-AF647^+^ neurons and CTB-AF555^+^ neurons along the mediolateral axis of the cortex of control and PAE mice. Data are mean ± SEM, n= 9 control mice from 3 litters, 18 sections, and, n= 9 PAE mice from 3 litters, 18 sections. CTB-AF647^+^ cells in bin 4 and 5: PAE vs control, p<0.0001 and p<0.01 respectively (Two-way ANOVA with Sidak’s multiple comparisons test). CTB-AF555^+^ cells in bin 11: PAE vs control, p<0.0001 (Two-way ANOVA with Sidak’s multiple comparison test). **C,** Images show retrogradely labelled neuronal cells bodies in the contralateral cortex. Arrowheads point to CPNs retrolabelled from the contralateral S1 area (green) in the M1 cortex of PAE mice. **D,** Fall latency of P60 control and PAE mice in the accelerated rotarod test. Data show the average of 3 daily trials ± SEM, n=30 control and 30 PAE mice from 3 litters; p= 0.8308 (Mann-Whitney test). **E,** Experimental design of the wheel running test. **F** and **G,** Wheel running performance of control and PAE mice tested at P60. Left: training on standard wheel, middle: complex wheels, right: complex wheels and treatment with tamapin. Data are mean ±SEM, n= 14 control and 15 PAE mice from 3 litters. Vmax day 8-10 (standard wheel) versus day 15-17 (complex wheel): p<0.0001 in PAE mice and p>0.05 in controls (Two-way ANOVA). Vmax day 15-17 (complex wheel) versus day 22-24 (complex wheel): p> 0.05 in PAE mice and p> 0.05 in controls (Two-way ANOVA). Scale bar: 500 µm (C).

### Defective motor skills in adult PAE mice

We then assessed whether the phenotype observed in the interhemispheric connectivity of adult PAE mice has an impact on behaviour. For this aim, we used a complex bimanual motor task in which mice run on a wheel with irregularly spaced rungs (‘complex’ wheel). This task engages motor control circuits and interhemispheric connecting pathways, and has previously detected behavioural deficits of CC transection or demyelination (45,46). We first showed that adult (P60) PAE mice did not show significant alterations in motor coordination in the accelerated rotarod test (Fig. 3D). The same cohorts of mice were used to assess their performance on running wheels following the schedule shown in Fig. 3E. PAE and control mice where first trained with a regular wheel with equally spaced rungs to get accustomed to the task. Both cohorts ran on the wheel spontaneously, spent the same amount of time turning the wheel, and improved their performance substantially over days, similarly increasing their daily distance and maximum speed (Vmax) attained (Fig. 3F and 3G). The results confirmed the absence of a general locomotor and learning defect in PAE mice. However, differences appeared between the groups when switching to the complex wheel. All mice experience difficulty at first, as shown by a drop in daily Vmax, but after a short 3-day adaptation period control mice could again run as fast on the complex wheel as on the normal wheel. In contrast, PAE mice failed to improve their speed, and their Vmax on complex wheels remained significantly lower compared to standard wheels throughout the week of testing (Fig. 3G). This was unlikely to result from a lack of motivation, since total number of runs, average run duration and total distance covered were the same for both groups (Fig. 3F and Supplementary Fig. 6C and 6D). These results show that exposure to moderate doses of alcohol during embryonic development has lifelong consequences for complex bilateral sensorimotor coordination requiring cortical control and interhemispheric communication.

A previous study reported that blocking the calcium-activated potassium channel Kcnn2 (Potassium Calcium-Activated Channel Subfamily N Member 2) mitigated motor skill learning deficits induced by PAE at higher dose (4.0 g/kg body weight) (47). To test the contribution of Kcnn2 on motor skill deficits in our PAE paradigm, the Kcnn2 channel blocker tamapin was administrated by daily intraperitoneal injections to control and PAE mice, and running parameters on complex wheels were recorded for an additional week. However, tamapin did not improve the locomotor deficits of PAE mice, which failed to improve their Vmax on complex wheels (Fig. 3G). These results suggest that changes in interhemispheric structural connectivity, rather than altered neuronal physiology, may underlie the deficits in locomotor coordination observed in our PAE mouse model.

### miR-17-5p, a candidate to mediate PAE effect on CC development

We then sought to identify mechanisms that might mediate the effect of EtOH on callosal formation. Alcohol is known to affect various epigenetic mechanisms, particularly microRNA expression. To test the hypothesis that PAE-deregulated miRNAs may have a cell autonomous role in callosal axon development, we first used a bibliographic approach to identify candidate miRNAs. We performed a cross comparison of miRNAs published to be deregulated by PAE (48–54), regardless of the animal model or exposure paradigm, and miRNAs known to be expressed in the axon of mouse embryonic cortical neurons (55). Among the 56 published PAE-deregulated miRNAs, 23 were expressed in growing cortical axons in vitro (Supplementary Table 1). Interestingly, one of them, miR-17-5p, was previously identified as a miRNA enriched in distal axons and growth cones of cultures mice cortical neurons (55), and is one of the seven miRNAs consistently reported in every axonal miARN profiling (56), suggesting an important function in axon extension and guidance. We used TargetScan (57) to predict miR-17-5p potential targets. The TargetScan Context+ score was then combined with gene expression levels in developing CPNs (58) using the TargetExpress method (59) to prioritise predicted targets of miR-17-5p likely to be functional in CPNs. Subsequent KEGG (Kyoto Encyclopedia of Genes and Genomes) pathway analysis revealed an overrepresentation of genes associated with the GO (Gene Ontology) terms ‘Axon Guidance’ and ‘Semaphorin and Ephrin signalling’, two signalling pathways critical for CC development (28,41,42,60,61). We therefore selected miR-17-5p for further analysis.

We first studied miR-17-5p expression in developing cortex in vivo. In situ hybridization for miR-17-5p on cryostat brain section reveals the presence of miR-17-5p in the cortical plate at E16.5, including in callosal neurons that were co-immunolabelled with the Satb2 marker (Fig. 4A and 4B). In addition, miR-17-5p was also detected in the CC at the midline of P0 brains, with higher expression in the ventral half of the tract (Fig. 4B). We next investigated how miR-17-5p levels were affected by EtOH in our PAE model. To do so, total RNAs were extracted from the cortex and microdissected CC regions of EtOH exposed and control P0 brains (Fig. 4C). Quantitative real-time polymerase chain reaction (qRT-PCR) showed that the expression of both miR-17-5p and the passenger strand miR-17-3p was significantly decreased after EtOH challenge in both cortex and CC, as was miR-9, which suppression has been previously reported to contribute to EtOH teratology (62) (Fig. 4D and 4E). Thus, miR-17-5p is a miRNA expressed by developing CPNs and their axons, whose levels are downregulated by PAE.

**Figure 4:**
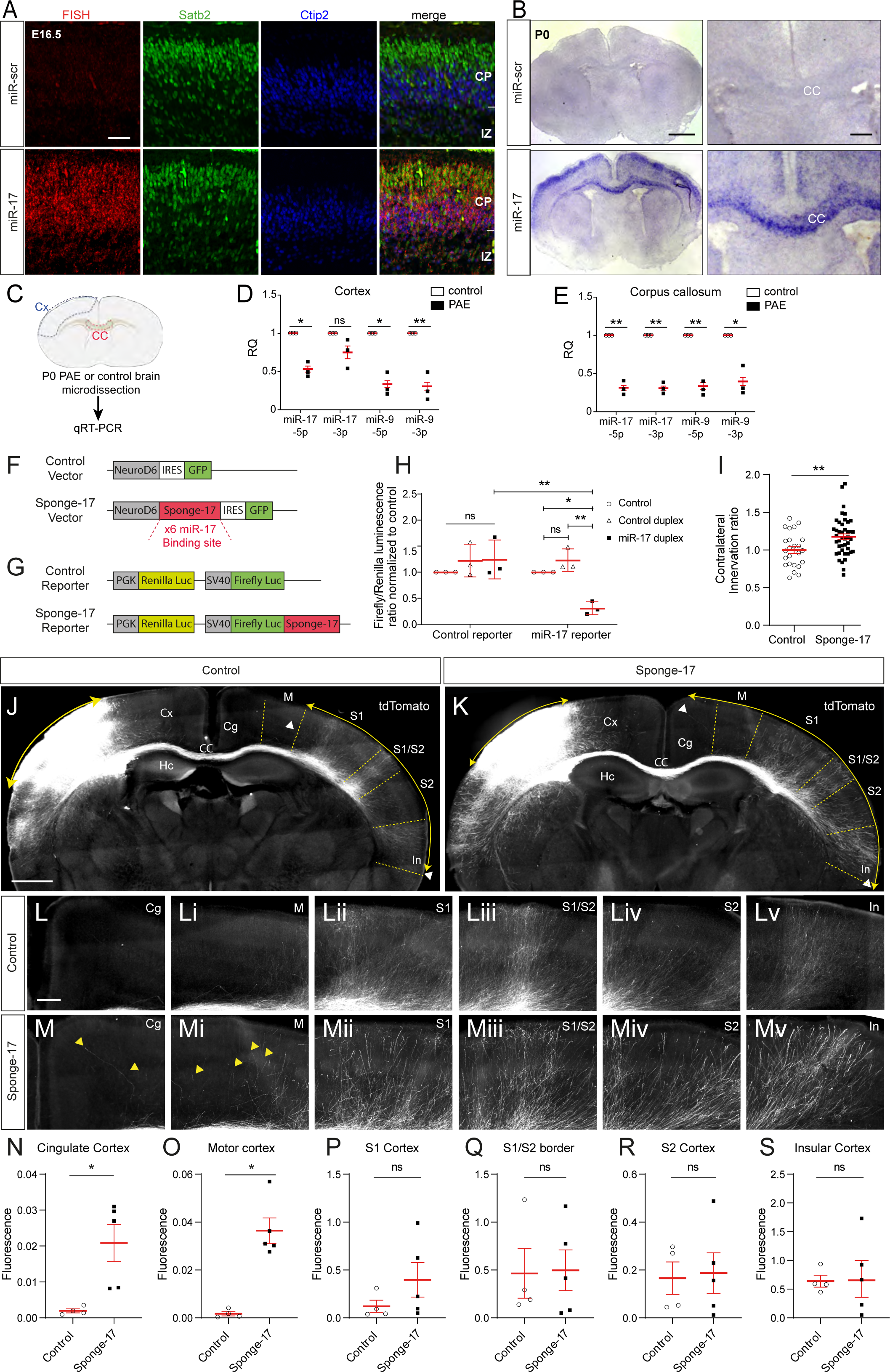
miR-17-5p is expressed in the developing CC, down-regulated by alcohol and its functional inhibition induces CC contralateral targeting defects. **A,** Fluorescent in situ hybridization (FISH) for miR-17-5p and negative control in section of E16.5 cortex combined with immunofluorescence staining for Satb2 and Ctip2. **B**, In situ hybridization detecting miR-17-5p in the P0 mouse brain. Negative control (non-targeting miR probes) from the same tissue is shown. **C**, Schematic showing brain regions dissected for qRT-PCR analysis. **D** and **E**, qRT-PCR analysis of expression of miR-17-5p, miR-17-3p, miR-9-5p and miR-9-3p in the cortex and CC of control and PAE mice at P0. n= 3 control and 3 PAE samples from 3 litters, 10 brains per sample. PAE vs controls: (D) miR-17-5p p= 0.0421, miR-17-3p p= 0.1232, miR-9-5p p= 0.01025, miR-9-3p p= 0.0099; (E) miR-17-5p p= 0.0023, miR-17-3p p= 0.0031, miR-9-5p p= 0.0085, miR-9-3p p= 0.0132 (Mann –Whitney test). **F** and **G**, Schematic of the miR-17-5p sponge (**F**) and luciferase reporter sensor vectors (**G**). **H**, Control and miR-17-5p luciferase reporter sensors were transfected into HEK 293T cells with miR-17-5p mimics or control duplexes. The relative luciferase expression ratio (firefly luciferase/Renilla luciferase) was calculated for three biological replicates. Data are mean ± SEM. Control reporter: control vs miR-17 duplex p> 0.05 (Two-way ANOVA, with Bonferroni’s multiple comparison); miR-17 reporter: control vs control duplex p>0.05, control vs miR-17 duplex p<0.05, control duplex vs miR-17 duplex p<0.01 (Two-way ANOVA, with Bonferroni’s multiple comparison); control reporter with miR-17 duplex vs miR-17-reporter with miR-17 duplex p= 0,0007 (Two-way ANOVA, with Sidak’s multiple comparison). **I,** Quantification of the contralateral innervation in brains electroporated at E15.5 with the miR-17-5p sponge (Sponge-17) or a control plasmid and examined at P8. Data are mean ± SEM, n=3 control mice from 2 litters, 24 sections, and n=5 PAE mice from 2 litters, 43 sections, p= 0.0087 (Mann Whitney test). **J** and **K**, Sections of P8 brains showing layer 2/3 callosal neurons of the somatosensory cortex electroporated by in utero electroporation at E15.5 with a tdTomato expressing plasmid together with the miR-17-5p sponge (Sponge-17) construct or a control plasmid. tdTomato^+^ projections innervate the contralateral cortical hemisphere. **L** and **M,** Higher resolution images of the different cortical regions shown in (**J**) and (**K**), revealing ectopic callosal projections in the cingulate and motor cortex of brains electroporated with the sponge construct (yellow arrowheads). **N-S**, Quantification of tdTomato fluorescence in the different cortical areas of the control and sponge-electroporated brains. Data are mean ± SEM, **N-S**, n=4 control mice from 2 litters, 4 sections, and n=5 PAE mice from 2 litters, 5 sections. **N**, p= 0.0159 (Mann Whitney test); **O**, p= 0.0159 (Mann Whitney test); **P**, p= 0.1905 (Mann Whitney test); **Q**, p= 0.9048 (Mann Whitney test); **R**, p= 0.9048 (Mann Whitney test); **S**, p= 0.9048 (Mann Whitney test). CC, corpus callosum; Cg, cingulate cortex; Cx, cortex; Hc, hippocampus, In, insular cortex; M, motor cortex; S1, primary somatosensory cortex; S2, secondary somatosensory cortex. Scale bars: 50 µm (A), 1000 µm (B, left panel), 250 µm (B, right panel).1200 µm (J,K), 300 µm (L,M)

### Functional inhibition of miR-17-5p phenocopies PAE-induced defects in CC development

To assess the function of miR-17-5p in developing CPNs, we used a functional knockdown approach using a miRNA sponge directed against miR-17-5p. The sponge was designed to contain six bulged (i.e., with 4 bases of mismatched) miR-17-5p binding sites (Fig. 4F) and act by titrating miR-17-5p from its endogenous targets. The efficacy of the sponge was validated using a dual-luciferase reporter assay. For this, we constructed a reporter sensor vector that encodes both a firefly luciferase reporter gene controlled by the miRNA-17-5p sponge sequence in its 3′ UTR and a Renilla luciferase normalization gene (Fig. 4G). The plasmid was transfected into HEK293T cells, along with mouse miR-17-5p mimic oligonucleotides or control mimics. Expression of miR-17-5p mimics, but not control mimics, induced a drastic decrease of luciferase activity (Fig. 4H). In contrast, transfection of miR-17-5p mimics had no effect on luciferase activity in HEK293T cells co-transfected with a control sensor plasmid lacking miR-17-5p binding sites (Fig. 4H). These data show that the designed miR-17-5p sponge functionally binds mouse miR-17-5p.

We next assessed the impact of a functional knockdown of miR-17-5p on CC development. To do so, the miR-17-5p sponge sequence was cloned under the control of the neurogenic differentiation 6 (NeuroD6) promoter in order to drive its expression in postmitotic neurons. The sponge construct or a control (empty) construct was delivered together with a tdTomato fluorescent reporter plasmid to developing CPNs by *in utero* electroporation of the somatosensory cortex at E15.5. Contralateral projections were first analysed at postnatal stages P3, when callosal axons had crossed the midline and grow within the intermediate zone of the contralateral cortical hemisphere (Supplementary Fig. 7A). Measurement of the distance between the axon growth front and the midline showed no significant difference between axons that received the sponge inhibitor and controls (Supplementary Fig. 7B and 7C), indicating that miR-17-5p does not regulate the growth rate of callosal axons. In contrast with the phenotype observed in PAE mice, S1 labelled axons that received the sponge remained in the ventral part of the CC. However, measurement of the distribution of labelled axons along the dorso-ventral axis (either pre-crossing, at the midline, and post-crossing) revealed a significant enlargement in the distribution of miR-17-5p sponge-electroporated axons compared to controls (Supplementary Fig. 7E). While this phenotype is milder than the one observed in PAE mice, this result still suggests that miR-17-5p is involved in the proper positioning of CC axons within the tract.

We next examined the contralateral targeting at P8. Data indicated that CPNs electroporated with the sponge inhibitor projected into a larger target area in the contralateral cortex compared with controls (Figure 4I-K). In addition, ectopic fibers entered the cingulate and motor cortices in mice electroporated with the miR-17-5p sponge inhibitor (Fig. 4J-4S). Overall, our results indicate that miR-17-5p is required for proper guidance and targeting of callosal axons, and that functional knockdown of miR-17-5p substantially phenocopies the effects of PAE on CC development.

### miR-17-5p targets in developing cortical neurons

To gain insight into the molecular mechanism by which miR-17-5p inhibition affects CC development, we searched for miR-17-5p regulated downstream gene in callosal neurons. To do so, we electroporated upper layer cortical neurons at E15.5 with the miR-17-5p sponge or a control construct, together with a tdTomato reporter, and isolated tdTomato^+^ neurons by fluorescence activated cell sorting (FACS) of dissociated cortices at P0 (Fig. 5A and Supplementary Fig. 8). We then extracted whole RNA from isolated neurons and performed RNA sequencing of each population (Supplementary Fig. 9).

**Figure 5.**
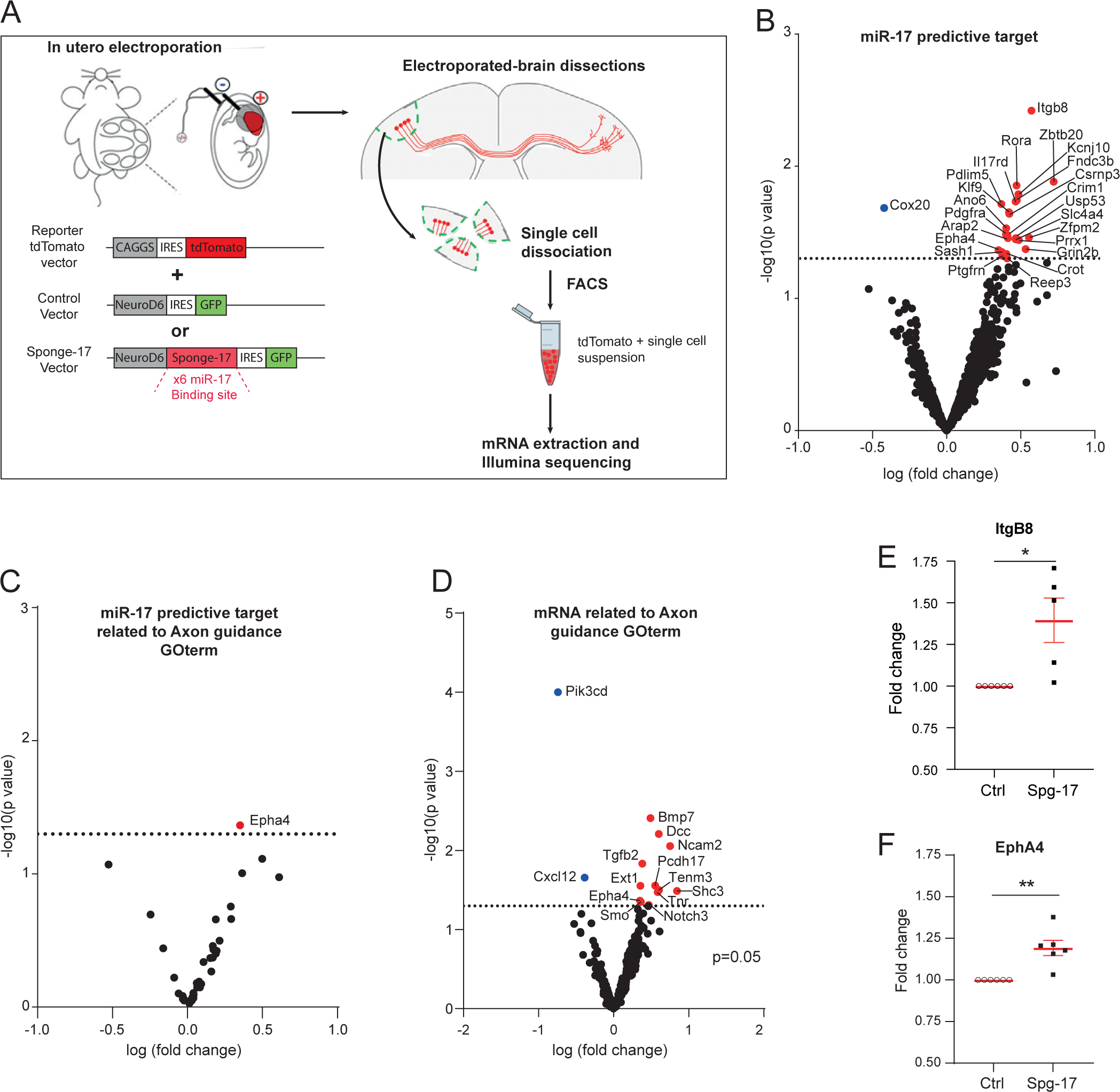
miR-17-5p sponge deregulated mRNA related to axon guidance GOterm. **A**. Illustration of experimental paradigm. **B**. Volcano plot of miR-17-5p predictive target deregulated in cortical neurons electroporated with the miR-17-5p sponge (Sponge-17). P value threshold set at 0.05. **C**. Volcano plot of miR-17-5p predictive target related to axon guidance GOterm deregulated in Sponge-17 condition. P value threshold set at 0.05. **D**. Volcano plot of mRNA related to axon guidance GOterm deregulated in Sponge-17 condition. P value threshold set at 0.05. **E, F.** Quantification of the relative expression levels of *Epha4* (**E**) and *Itgb8* (**F**) mRNAs by RT-qPCR in cortical neurons electroporated with a control plasmid or the miR17-5p sponge (Spg-17) plasmid. Data are mean ± SEM, n = 6 control and PAE mice from 3 litter; **E,** p= 0.0082 (one sample t-test); **F,** p= 0,0414 (one sample t-test).

Among the 10,812 genes detected, 822 were bioinformatically predicted to be targeted by miR-17-5p (TargetScan) (57). To gain insight into the biological processes associated with these targets we performed a gene ontology (GO) analysis based on biological process GO terms and KEGG pathways. « Axon Guidance » pathway was found to be the most significantly enriched with an FDR enrichment score of 2.24*10^-10^, suggesting that miR-17-5p might directly regulate axon guidance signaling in CPNs. Comparison of sequencing data between neurons in which miR-17-5p was inhibited and controls identified a total of 222 mRNAs that were differentially expressed, including 24 miR-17-5p potential targets (23 upregulated and 1 downregulated) (Fig. 5B). Among them, one gene was associated to the Axon Guidance GOterm (GO:0007411), the upregulated axon guidance receptor Ephrin type-A receptor 4 (EphA4) (Fig. 5C). Interestingly, EphA4 is expressed in developing CC neurons (63) and has been validated as a direct miR-17-5p target in heterologous cells (ref.).

Differential expression of *Epha4* between PAE and control samples was confirmed by quantitative real-time PCR (qPCR) (Fig. 5F and 5E). In addition, 198 genes not predicted to be targeted by miR-17-5p were found to be deregulated in neurons in which miR-17-5p was inhibited, including 13 genes associated to the Axon Guidance GOterm (11 upregulated and 2 downregulated) (Fig. 5D). Among those genes were : the axon guidance receptor deleted in colorectal cancer (DCC), which is expressed in developing CC axons (64), the glycoprotein Teneurin-3 (Tenm3), which regulates the topographic patterning of diverse axonal projections (65–67) and the transmembrane glycosyltransferase exostosin 1 (Ext1), which is required for the proper growth and guidance of CC axons (68). Together, these results suggest that miR-17-5p has the capacity to alter the expression of several axon guidance genes either by directly or indirectly interacting with downstream target transcripts.

### Overexpression of EphA4 phenocopies PAE-induced defects in CC development

The above results suggest that PAE-induced suppression of miR-17-5p could lead to EphA4 upregulation in CC neurons, thereby contributing to CC development defects. Indeed, Eph signaling has been reported to regulate the segregation of medial and lateral cortical axons within the CC (41) and the formation of topographic map in the nervous system (69). To test this hypothesis, we first evaluated the expression level of EphA4 in callosal neurons after PAE. To do so, we electroporated upper layer cortical neurons at E15.5 with a tdTomato reporter, challenged the embryos with EtOH or vehicle, and isolated tdTomato^+^ neurons by FACS of dissociated cortices at P0. PAE caused a significant increase in the expression level of *Epha4* measured by qPCR, and the same trend was seen for *Itgb8*, although not statistically significant (Fig. 6A and 6B). We next analyzed the expression pattern of EphA4 in the CC of control of PAE mice by immunostaining at P6. In accordance to previously published data (70), EphA4 signal was restricted to the most dorsal CC fibers in controls (Fig. 6C). This pattern inversely correlated with the pattern of expression observed for miR-17-5p (Fig. 4B). In the CC of PAE mice, the EphA4 signal was increased in the more ventral parts of the CC (Fig. 6C-6E). Overall, our results suggest that PAE-induced miR-17-5p knockdown results in upregulation of EphA4 in ventral CC axons, which normally express higher levels of miR-17-5p.

**Figure 6.**
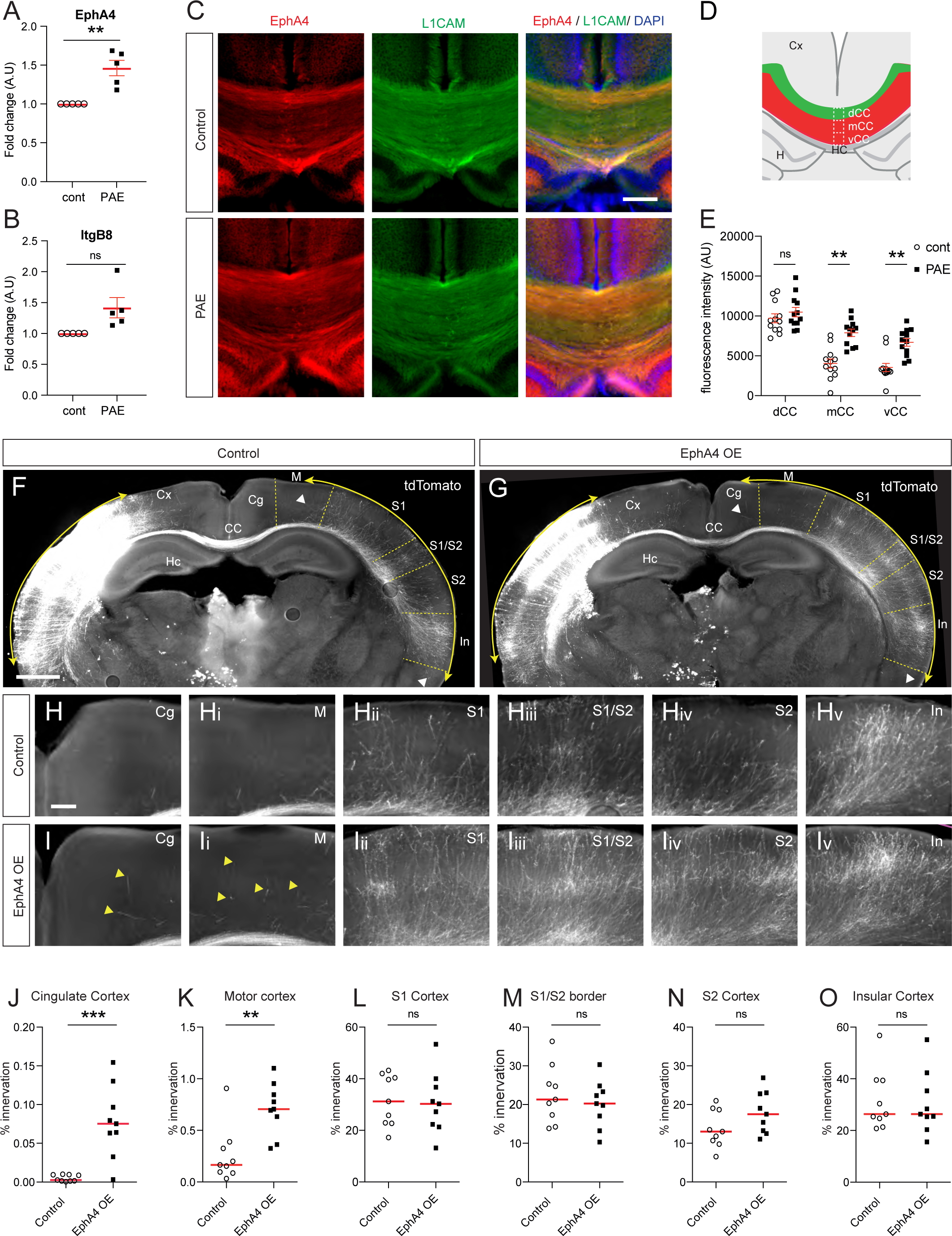
PAE upregulates EphA4 expression and EphA4 overexpression induces contralateral targeting defects. **A** and **B,** Quantification of the relative expression levels of *Epha4* and *Itgb8* by RT-qPCR in electroporated cortical neurons from control and PAE mice at P0. Data are mean ± SEM, n=5 control and PAE mice from 3 litter; **A**, p= 0,0097 (one sample t-test); **B**, p= 0,0644 (one sample t-test). **C**, Immunostaining of L1CAM and EphA4 on tissue sections through the corpus callosum of PAE and control mice at P6. **D,** Schematic illustrating measurement methods for analysis of EphA4 expression pattern in the CC. **E**, Quantification of EphA4 immunostaining in the dorsal, medial and ventral CC on sections from control and PAE mice at P6. Data are mean ±SEM, n= 6 cont mice from 2 litters, 12 sections and n=5 PAE mice from 2 litters, 12 sections. dCC control vs PAE, p= 0.5185 ; mCC control vs PAE p=0.0014 ; vCC control vs PAE p=0.0024 (Multiple Wilcoxon Tests). **F** and **G**, Sections of P8 brains showing layer 2/3 callosal neurons of the somatosensory cortex electroporated by in utero electroporation at E15.5 with a tdTomato expressing plasmid together with control or EphA4-expressing plasmids. tdTomato^+^ projections innervate the contralateral cortical hemisphere. **H** and **I,** Higher resolution images of the different cortical regions shown in (F) and (G), revealing ectopic callosal projections in the cingulate and motor cortex of brains electroporated with the EphA4 plasmid (yellow arrowheads). **J-O**, Quantification of tdTomato fluorescence in the different cortical areas of the control and sponge-electroporated brains. Data are mean ± SEM, n = 5 control and 5 EPHA4 OE mice, from 2 litters, 9 sections. **J**, p= 0.0005 (Mann Whitney test); **K**, p= 0.0056 (Mann Whitney test); **L**, p= 0.4894 (Mann Whitney test); **M**, p= 0.6665 (Mann Whitney test); **N**, p= 0.1359 (Mann Whitney test); **O**, p= 0.9314 (Mann Whitney test). CC, corpus callosum; Cg, cingulate cortex; Cx, cortex; Hc, hippocampus, In, insular cortex; M, motor cortex; S1, primary somatosensory cortex; S2, secondary somatosensory cortex. Scale bars: 200 µm (C), 1200 µm (F, G), 300 µm (H, I).

We next assessed the impact of an overexpression of EphA4 in callosal neurons projecting through the ventral CC, i.e callosal neurons from the somatosensory cortex. To do so, a mouse *Epha4* expressing plasmid construct was delivered together with a tdTomato fluorescent reporter plasmid to developing CPNs by *in utero* electroporation of the somatosensory cortex at E15.5. Contralateral targeting was analysed at postnatal stage P8. Data indicated that CPNs electroporated with the *Epha4* plasmid projected to a larger target area in the contralateral cortex (Figure 6F-6G) and ectopically innervated the cingulate and motor cortices (Fig. 6H-O). Overall, our results indicate that proper expression of EphA4 is required for contralateral CC axon targeting and that overexpression of EphA4 phenocopies targeting defects induced by miR-17-5p functional knockdown and PAE.

## Discussion

PAE is known to affect brain development and function, leading to a wide range of neurological conditions. However, while brain connectivity is affected, either structurally or functionally, in FASD patient, it is not known whether PAE directly affects the ability of neuronal axons to extend and be guided to their target. Here we report that PAE causes defects in CC development, but not in neural guidepost structures, strongly suggesting that alcohol alters the intrinsic axonal mechanisms governing growth and guidance. This hypothesis is further supported by the finding of the key role for miR-17-5p-mediated repression of *Epha4* mRNA in CPNs in establishing interhemispheric connectivity. Exposure to alcohol during development suppresses miR-17-5p expression in CPNs, such that the sharp expression domain of EphA4 in the dorsal part of the CC extends to the ventral domain. This ectopic overexpression of EphA4 results in abnormal topographical segregation of callosal fibers within the CC and targeting errors in the opposite cortical hemisphere, which persist into adulthood. These results highlight axon-intrinsic molecular pathways whose deregulation by alcohol affects the structural organisation of neural circuits and may provide the neurological substrate for behavioural alterations in FASD.

Despite the similarity between the brain wiring phenotypes induced by PAE, miR-17-5p knockdown and EphA4 overexpression, the deregulation of the miR-17-5p/*Epha4* pair is not sufficient to explain the full range of axonal developmental defects induced by PAE. For example, the axonal growth delay that may contribute to CC hypoplasia at birth was not reproduced by functional knockdown of miR-17-5p, suggesting the involvement of other mechanisms. Interestingly, miR-17-5p belongs to the polycistronic group miR-17-92, which encodes six individual, frequently co-expressed miRNAs that target genes enriched in the same biological pathways. Among the miR-17-92 cluster, miR-19a is a key mediator of axonal growth in embryonic cortical neurons though local modulation of PTEN (phosphatase tensin homolog) protein levels (71). Thus, the previously reported downregulation of miR-19a by PAE (48) could contribute to the observed delay in callosal axon growth. On the other hand, a large body of literature has shown that alcohol can directly bind to the extracellular domain of the L1CAM protein, which is richly expressed by CC fibers, and inhibit L1CAM-mediated cell adhesion processes such as neurite outgrowth (72,73). It is therefore likely that the effects of PAEs on the development of callosal projections result from the deregulation of multiple molecular pathways.

Our results demonstrate defects in the areal targeting of callosal axons in the contralateral cortex, a process that occurs one week after the end of alcohol exposure in our EAP protocol. How can the influence of alcohol persist in later stages of CC development? A first hypothesis is based on the finding that fiber-fiber interactions in the tract and/or the dorso-ventral position at which callosal axons cross the brain midline determine their subsequent projection site in the contralateral cortex (74). Hence, aberrant targeting of callosal fibers in PAE mice could be a direct consequence of prenatal alcohol-induced alterations at the CC midline. Another possibility is that EAP may cause a persistent change in the cell-intrinsic mechanisms underlying axon navigation. Indeed, recent studies have shown that PAEs cause long-lasting changes in neuronal epigenetic marks and gene expression, leaving a ‘memory’ of exposure (47). In this context, it is possible that persistent changes in miR-17-5p and EphA4 expression after the end of alcohol treatment subsequently disrupt axon-target interactions that regulate the wiring pattern of callosal projections.

Our data showed an alcohol-induced reduction in miR-17-5p levels, but the underlying mechanisms are not elucidated. In general, alcohol metabolism is known to altering the epigenetic landscape via modulation of DNA/histone acetylation and methylation of specific set of genes and miRNA loci (75–81). Increased acetylation triggers feedback mechanisms, such as overexpression of histone deacetylases (HDACs), designed to restore basal levels of histone acetylation (82,83). Interestingly, the miR-17-92 cluster has been reported to be directly represses by HDAC3 and HDAC9 (82,84–87), suggesting HDAC expression/activation as one of the possible mechanisms to control alcohol-induced suppression of miR-17-5p. In addition, alcohol metabolism induces important endoplasmic reticulum (ER) stress (88). Interestingly, diverse ER-stress sensors such as the inositol-requiering enzyme 1 IRE1 and the PRKR-like endoplasmic reticulum kinase PERK have been shown to negatively regulate miR-17-5p (89,90). Thus, an ER Stress-mediated response could be a potential mechanism for the altered miR-17-5p expression after PAE, a hypothesis that remains to be explored.

Finally, we have shown here that in utero exposure to moderate doses of alcohol has lifelong consequences on the behaviour of mice measured in a complex bilateral sensorimotor coordination task previously used to reveal deficits induced by CC transection or demyelination (45,46). Earlier work has shown that prenatal exposure to higher doses of alcohol than those used in this study (4 g/kg instead of 2g/kg) induces motor learning deficits in adult mice by altering the expression of the calcium-dependent potassium channel Kcnn2 and the physiological properties of motor cortical neurons (47). Here we show and that the observed deficits in complex locomotor coordination cannot be rescued by administration of the Kcnn2 channel blocker tamapin. This strongly suggests that alterations in structural connectivity between the cortical hemispheres, rather than functional defects, are responsible for the complex bilateral sensorimotor coordination defect in the present EAP mouse model.

It should be noted that behavioural defects in PAE mice were only detected when the animals were subjected to complex tasks (i.e., running on a wheel with irregularly spaced rungs). Similarly, in humans, children with non-severe FASD do not show behavioural or cognitive impairments on simple tasks, but experience difficulties when tasks become more complex (91,92). Interestingly, the asymmetry of cortical functional organisation of the human brain implies that interhemispheric information transfer increases with task complexity (93). This suggests that mild defects in interhemispheric connectivity such as those we observed in PEA mice may have more detrimental consequences for cognitive and motor performances in humans.

## Acknowledgments

The authors want to acknowledge the collaboration with Dr Gregor Kasprian and his team at the Medical University of Vienna, Department of Biomedical Imaging and Image-guided Therapy. We also thank Christophe Beclin (IBDM), Andrea Erni (IBDM) and Micaela Roque (IBDM) for their advises and substantial help with the experiments. This work was supported by Centre National de la Recherche Scientifique (CNRS), France ; Aix Marseille Université, France ; Institut National de la Santé et de la Recherche Médicale (INSERM), France ; grants from ANR (WIRING-FASDBRAIN, ANR-18-CE91-0009) to F.M. The France-BioImaging infrastructure is supported by the Agence Nationale de la Recherche (ANR-10-INBS-04-01, ‘Investissements d’Avenir’). M.A. is a recipient of a doctoral fellowship from the French Ministry of Higher Education, Research and Innovation (MESRI).

## Author Contributions

Acquisition of data: M.A., A.B.. Analysis and interpretation of data: M.A., A.B., F.M. Study supervision : A.B., F.M.

## Supplementary methods

### Dual luciferase assay

To validate the sponge, the sequence containing 6MBS for miR-17 was inserted into pmirGLO dual luciferase miRNA target expression vector (Cat. #E1330; Promega) downstream of the Firefly luciferase using XbaI and SalI. HEK293T cells were transfected with a Lipofectamine 3000 Kit (Cat. # L3000001; ThermoFisher Scientific). 0.7 µg of plasmid (control reporter or miR17-5p sponge reporter) and 50 pmol of miR-17 or negative control miRNA mimic (miRIDIAN microRNA mimic, miR-cont Cat. #CN-001000-01-05, miR-17 Cat. # C-310561-07-0002; Dharmacon^TM^) were used for lipofection. The luciferase assay was performed with a Dual Luciferase Assay kit (Cat. # E1960, Promega) and a Mithras LB 940 luminometer (Berthold, Bad Wildbad, Germany) according to manufacturer instructions after 36-48.

### In situ hybridization on cryosection

ISH were performed on 16 µm-thick cryosections. miR-17 LNA probes (Cat. #339111; Qiagen) or control probes were used at a concentration of 10 nM for colorimetric labeling and 20 nM for fluorescent labeling. The colorimetric revelation was performed using alkaline phosphatase (AP) coupled anti-DIG antibody (1/2000, Cat. #11093274910; Sigma) and NBT-BCIP. For immunostainong and fluorescent revelation, sections were blocked in 0.5 % blocking agent (Cat. # FP1020; Perkin Elmer) and incubated with anti-DIG POD antibody (1/2000, Cat. #11207733910; Sigma) and antibodies against Satb2 and Ctip2 (see Supplementary Table 3). Stainings were revealed with TSA Plus Cy3 Kit (Cat. # NEL744001KT; Perkin Elmer) and secondary antibodies (see Supplementary Table 3). Cell nuclei were stained with NucBlue Fixed Cell ReadyProbes reagent (Cat. #R37606; ThermoFisher Scientific, Waltham, USA).

### Wholemount in situ hybridization

Whole brains were dissected out and meninges were removed prior fixation. ISH was performed with 200ng/mL of BHLHB5 or LMO4 probes. Revelation was performed with anti-DIG-AP antibody (1/4000, Cat. #11093274910; Sigma) and NBT BCIP solution (1X ready to use solution, Roche).

### Fluorescence immunohistochemistry

80-100 µm thick Vibratome brain sections (80-150 µm-thick) were immunolabeled with primary antibodies and secondary antibodies, listed in Supplementary Table 3, diluted in 0.3 % Triton X-100 and 10 % donkey serum in PBS. Cell nuclei were stained with NucBlue Fixed Cell ReadyProbes reagent (Cat. #R37606; ThermoFisher Scientific, Waltham, USA).

### Whole brain immunostaining and clearing

P3 brains were fixed for 4 h in 4% PFA in PBS, and then immunolabeled and cleared using a modification from the iDISCO+ protocol (94). The blocking solution from the iDISCO+ protocol was replaced by a solution of 0.2 % gelatin, 0.1 % saponin, 0.5 % Triton X-100, 0.01 % Thiomersal (Cat. # PHR1587-1G; Sigma) in PBS.

### Sequencing and analysis

Librairies were generated using Illumina Stranded mRNA Prep with poly(A) selection and sequenced to at least 120 million mapped paired-end reads per replicate (Supplemental figure 8D). 73bp paired-end reads were mapped to the mouse genome (mm10) using STAR (2.7.6a) with an average of 1.24 ×10^8^ mapped reads (range 1.15×10^8^-1.38×10^8^; s.d. 8.45×10^6^). Mapped reads from individual replicates were assembled into transcripts using Samtools (1.11) with default parameters. Differential expression was obtained with DESeq2 (v1.30.1) that normalized data by relative log expression. Data normalization have almost no effect on data distribution (Supplemental figure 8D). A threshold of 1000 count was applied before further analysis. miR-17 potential targets were identified on TargetScan v 7.2 (57) and Gene ontology (GO) enrichment analysis was performed using ShinyGO v0.74 (95). GO terms and terms associated with KEGG pathways were considered if their enrichment false discovery rate *p* value was ≤0.05.

### Layering distribution

To quantify the layer position of labelled cells, we divided the cortical thickness into 10 bins (96). In S1, bins 2–3 represent layers 2/3, bin 4–5 represent layer 4, bin 6 represents layer 5A, bin 7 represents layer 5B, and bins 8–10 represent layer 6. In M1, bins 2–4 represent layers 2/3, bin 5 is layer 5A, bins 6–7 are layer 5B, and bins 8–10 are layer 6. Cell counting was performed using the Photoshop counting tool.

### Image acquisition

Fluorescent images of vibratome and cryostat brain sections were acquired with an Apotome imager M2 (ZEISS) equipped with a Hamamatsu C11440 digital camera, or a Confocal LSM880 (ZEISS) equipped with PMT, CH3, and GAasp photomultipliers detectors. Colorimetric staining was imaged with an AxioImager M2 (ZEISS) equipped with a CamColor camera, or with a LUMAR V12 Stereoscope (ZEISS) equipped with a CamColor camera when a larger acquisition field was required. ZEN 2012 acquisition software was used for image acquisition on the Apotome and Confocal microscopes, and AxioVision software was used for image acquisition with the AxioImager and the LUMAR. The images were transformed into Tiff and processed and quantified on FIJI (ImageJ) and Adobe Photoshop 2019. Cleared brains were imaged with a light-sheet fluorescence microscope (Ultramicroscope II; LaVision BioTec) equipped with the ImSpector pro acquisition software. The sample and the objective were immersed in a tank filled with dibenzyl ether to maintain a homogeneous refractive index between the specimen and the objective. The sample was imaged in the coronal direction and an image was acquired every 2.5 µm, with a bilateral illumination and a maximum gray level between 20,000 and 60,000 depending on the staining quality. Images were exported to the 3D image processing software Imaris (Bitplane) to perform 3D reconstructions and then process the images.

### Dorso ventral distribution of Electroporated axon in the CC

The dorso-ventral distribution of electroporated axons was quantified on one coronal optical slice of clarified and 3D reconstructed brains at the level of the Septum, fornix and hippocampal commissure.

### miRNA RT-qPCR

Regions corresponding to the CC and premotor/ presensory cortex were manually dissected from 150um thick vibratome sections (Leica VT1000S) of unfixed brains from control or PAE P0 individuals. Total RNA (10 brains per condition) was extracted with Trizol (Cat. #15596018; Invitrogene), miRNAs were retrotranscribed using miRCURY LNA RT kit (Cat. #339340) and qRT-PCR performed using miRCURY LNA SYBR® Green PCR Kit (Cat. # 339345). Reactions were run on a Bio-Rad CFX96 Real-Time System. For quantitative analysis, cycle threshold (Ct) mean values were measured in technical and biological triplicates, and the ΔΔCt method (97) was applied using U6 as normalizing control. The following miRCURY LNA probe were used: has-miR-17-5p, (Cat. # YP02119304; Qiagen), hsa-miR-17-3p, (Cat. # YP00205325; Qiagen), hsa-miR-9-5p, (Cat. # YP00204513; Qiagen), has-miR-9-3p, (Cat. # YP00204620; Qiagen), and the normalizing control (hashsa, Cat. # YP00203807; Qiagen)

### RT-qPCR

Total RNA of FACS-sorted tdTomato cell suspension coming from, miR-17 sponge electroporated, ETOH-exposed, or control P0 brain were extract with Trizol and were retrotranscribed using SuperScript III Reverse Transcriptase (Cat. #18080-044, Invitrogen). qRT-PCR were performed using PowerUp SYBR® Green master mix (Cat. # A25742, Thermofisher scientific). Reactions were run on a Bio-Rad CFX96 Real-Time System. For quantitative analysis, cycle threshold (Ct) mean values were measured in technical and biological triplicates, and the ΔΔCt method (97) was applied using β-ACT as normalizing control. The following PCR primer were used: EPHA4 F-TGGAATTTGCGACGCTGTCA and R-CACTTCCTCCCACCCTCCTT, ITGβ8 F-ATGCTTCAGGCTGCCGTCTGTG and R-CAGTTTCCGTCATTCGGCACCA, and β-ACT F-GCTGTGCTGTCCCTGTATGCCTCT and R-CCTCTCAGCTGTGGTGGTGAAGC.

### Statistics

Statistical analyses were performed using Graphpad Prism version 6 or 9 (GraphPad Software Inc., La Jolla, CA). Normal distribution of the data was examined using the D’Agostino–Pearson omnibus, Shapiro-Wilk, or Kolmagorov-Smirnov tests. Statistical test used are described in figure legends.

